# Microbiome Dynamics During Translocation of the Critically-Endangered Frog *Craugastor ranoides*

**DOI:** 10.1101/2025.08.06.668623

**Authors:** Xavier A Harrison, Katherine Herborn, Gilbert Alvarado, Maria Marta Chavarría, Kirsty J Marsh, Alexander Wilson, Alexandra North, Victoria EP Wakeham, Roberto Fernández, Hector Zumbado, Jon Bielby, Robert Puschendorf

**Affiliations:** Centre for Ecology & Conservation, University of Exeter, UK; School of Biological and Marine Sciences, University of Plymouth, UK; Escuela de Biología, Universidad de Costa Rica, San Pedro, Costa Rica; Programa de investigación, Área de Conservación Guanacaste, Liberia, Costa Rica; Guanacaste Dry Forest Conservation Fund, Philadelphia, USA; Escuela de Ciencias Biologicas, Universidad Nacional de Costa Rica, Heredia, Costa Rica; School of Biological and Environmental Sciences, Liverpool John Moores University, Liverpool, UK

**Keywords:** skin bacterial community, microbiota, conservation translocation, amphibian declines, *Batrachochytrium dendrobatidis*

## Abstract

1. Amphibian populations are declining and species going extinct due to habitat loss, climate change and emerging infectious disease, prompting the increasing use of translocations as a conservation tool.
2. Skin-associated microbial communities play a key role in amphibian health, particularly in protecting hosts from pathogenic infection. However, to date we know relatively little about the impact of translocation on the dynamics of skin-associated microbes, or the consequences of translocation-mediated disturbance of the microbiome for amphibian health.
3. We investigated the skin microbiome of the critically endangered frog Craugastor ranoides, now restricted to a fraction of its historical range in Costa Rica. We first conducted a multi-population survey to characterise among-population variation in skin bacterial communities. We then translocated individuals to a new site, using radio tracking to monitor them over time and assess individual-level changes in body mass, microbiome composition, and predicted microbial function.
4. We found limited among-population variation in skin microbiome composition across three different populations of *C. ranoides*. Response of the microbiome to translocation was highly individual-specific; some frogs demonstrated relatively stable community richness and composition, whilst others exhibited marked turnover in bacterial membership and predicted function. Body mass showed similar dynamics, but several of the recaptured frogs had gained mass after several days in the translocation site.
5. *Synthesis and Applications*: Our findings highlight the importance of considering microbiome dynamics in threatened species translocations to optimize individual health. This includes considering whether individuals should be prioritised for translocation based on their initial microbiome composition, and understanding how shifts in microbiome post-translocation may alter overall success of translocation efforts. These insights extend beyond amphibians, offering a broader framework for integrating long-term monitoring of host microbiomes and health into conservation efforts.

## INTRODUCTION

Amphibians have experienced population collapses and even extinctions across the globe. Over 40% of species are threatened due to habitat loss, climate change, invasive species, and emerging infectious diseases (Lips et al., 2006; Scheele et al., 2019; Stuart et al., 2004). These declines are especially severe in tropical and high-elevation regions, where entire amphibian communities collapsed within a few years, often in pristine, protected habitats. For a long time, these were considered “enigmatic declines” until the emergence of chytridiomycosis was recognized as the primary driver (La Marca et al., 2005; Lips et al., 2006; Scheele et al., 2019). Habitat protection remains fundamental to amphibian conservation as some species that suffered enigmatic declines were first presumed extinct, but have since been rediscovered and recovered in places in which suitable habitat protection has been afforded (García-Rodríguez et al., 2012; Scheele et al., 2017). However, these small and relictual populations remain at significant risk of extinction due to their limited size, which increases vulnerability to environmental change, demographic stochasticity, and disease outbreaks (Smith et al., 2009).

Given the high risk of extinction faced by small, isolated populations due to stochastic events and natural population cycles, ex situ population management has increasingly been promoted as a means to ensure long-term species survival. Captive breeding, once the backbone to conservation of amphibians threatened by chytridiomycosis (Mendelson et al., 2006) can introduce its own challenges, including fitness loss associated with altered or diminished microbiomes (Ruthsatz et al., 2020). Translocation, moving individuals from captive or source populations to restored or safer habitats, has become an increasingly important conservation strategy for threatened amphibians. Given the critical role of skin-associated microbial communities in amphibian health (Bates et al., 2018; Harrison et al., 2019), understanding how microbiomes contribute to disease resistance and post-translocation survival is essential (Dallas and Warne, 2023a).

In amphibians, the skin microbiome is particularly important, forming the first line of defense against pathogens while also contributing to physiological processes such as osmoregulation and cutaneous respiration (Bates et al., 2022). Amid escalating disease threats and rapid climate change, research increasingly highlights the amphibian microbiome as a critical line of defense, with certain bacterial taxa producing antifungal and antiviral compounds that inhibit the growth of pathogenic organisms (Harrison et al., 2019; Rebollar et al., 2018). However, these microbial communities are not static; they are highly dynamic (Marsh et al., 2024), shifting in response to environmental factors such as temperature, humidity, and substrate composition, raising questions about the stability of host-microbe associations (Longo et al., 2015). These shifts in microbial composition due to environmental change or translocation may influence disease susceptibility (Buttimer et al., 2024). While some individuals retain core microbial taxa that provide consistent protection, others experience turnover that could either enhance or diminish their defenses (Loudon et al., 2014). This interplay between microbiome stability, disease risk, and environmental change is a critical but poorly understood factor in amphibian conservation, especially when implementing conservation based translocations (West et al., 2019). Understanding microbiome variation among populations is particularly important in conservation biology, as microbial composition may influence an individual’s resilience and fitness to new environmental pressures, including pathogen exposure (Kueneman et al., 2014).

Given the high costs and limitations of captive breeding, translocation has become a widely used conservation strategy for amphibians, particularly amid global population declines. However, amphibian translocation success remains variable—with reported success rates hovering around 50%, and failures often linked to small release numbers, poor site selection, and lack of long-term monitoring (Germano and Bishop, 2009; Hossack et al., 2022b). To maximize success, best practices recommend phased translocations with pilot studies to identify and mitigate these risks. In this context, assessing microbiome diversity across populations can offer valuable insights into whether individuals possess beneficial microbial communities that may enhance their survival in novel habitats (Kueneman et al., 2016).

The case of the critically endangered frog *Craugastor ranoides* highlights the urgency of integrating microbiome research into conservation efforts. Once widely distributed across lowland dry forests and high-elevation cloud forests in Costa Rica, this species suffered extensive declines during the global amphibian crisis, disappearing entirely from montane habitats (Puschendorf et al., 2006, 2005). The surviving populations now persist only in isolated tropical dry forest refugia from chytridiomycosis, where high temperatures and lower humidity may have limited pathogen prevalence (Puschendorf et al., 2009; Zumbado-Ulate et al., 2014). However, these habitats are now facing unprecedented environmental stress due to climate change. During the 2014-2015 El Niño event, the region experienced extreme drought, with rainfall falling to just 39% of the long-term mean in some areas, with a very narrow distribution, most of this in just a few weeks at the start of the wet season followed by a prolonged drought (Campos et al., 2020). This climatic anomaly triggered ecosystem-wide impacts, including a 42% decline in gross primary productivity (Castro et al., 2018), and widespread tree mortality driven by species-specific hydraulic failure. In the dry forest of the Área de Conservación Guanacaste (ACG), 13% of 1,577 individually monitored trees died, with some species such as *Quercus oleoides* experiencing up to 34% mortality (Powers et al., 2020). These changes in forest structure and canopy cover likely altered microhabitats critical for amphibian survival. At the same time, *C. ranoides* populations in the dry forest of ACG collapsed entirely following the complete drying of a breeding stream, and no individuals were observed at the site after the event (Puschendorf et al., 2019).

This convergence of climatic and ecological stressors requires urgent conservation intervention. Efforts are currently underway to test whether translocating individuals back to the volcanic cloud forests survive, where cooler and wetter conditions may provide a more stable environment. However, the microbiome’s role in this translocation remains unclear, particularly regarding whether individuals retain beneficial microbial communities that could support their survival in the montane environment. The aims of this study are to i) quantify among-population variation in skin bacterial community composition in *Craugastor. ranoides*; and ii) investigate the effect of translocation on *C. ranoides* skin microbiota dynamics and whether these changes relate to host health. We monitored microbiome composition in frogs before and after experimental translocation to premonate rainforest and compared them to individuals sampled on stream transects in their native dry forest habitat.

## METHODS

### Field sites and sampling

Fieldwork took place across four discrete periods: January 3–12, 2022; April 7–14, 2022; January 5–14, 2023; and March 9–16, 2023, with the final period coinciding with a translocation event.

Background sampling for microbiome characterization was conducted at Murciélago, Pedregal, and Brazo del Río Potrero Grande (Figure 1), Área de Conservacion Guanacaste, in the northwest of Costa Rica. Frogs were captured by hand while wearing nitrile gloves to prevent cross-contamination between individuals. Skin microbiome samples were collected by swabbing both the dorsal and ventral surfaces of each frog using MW100 DrySwabs (Medical Wire Equipment), and swab tips were stored in absolute ethanol at ambient temperature. Snout–vent length (SVL) was measured with digital calipers to the nearest millimeter, and mass was recorded using Pesola Light-Line Precision Spring Scales. All individuals were released at their exact point of capture following sampling.

**Figure 1.**
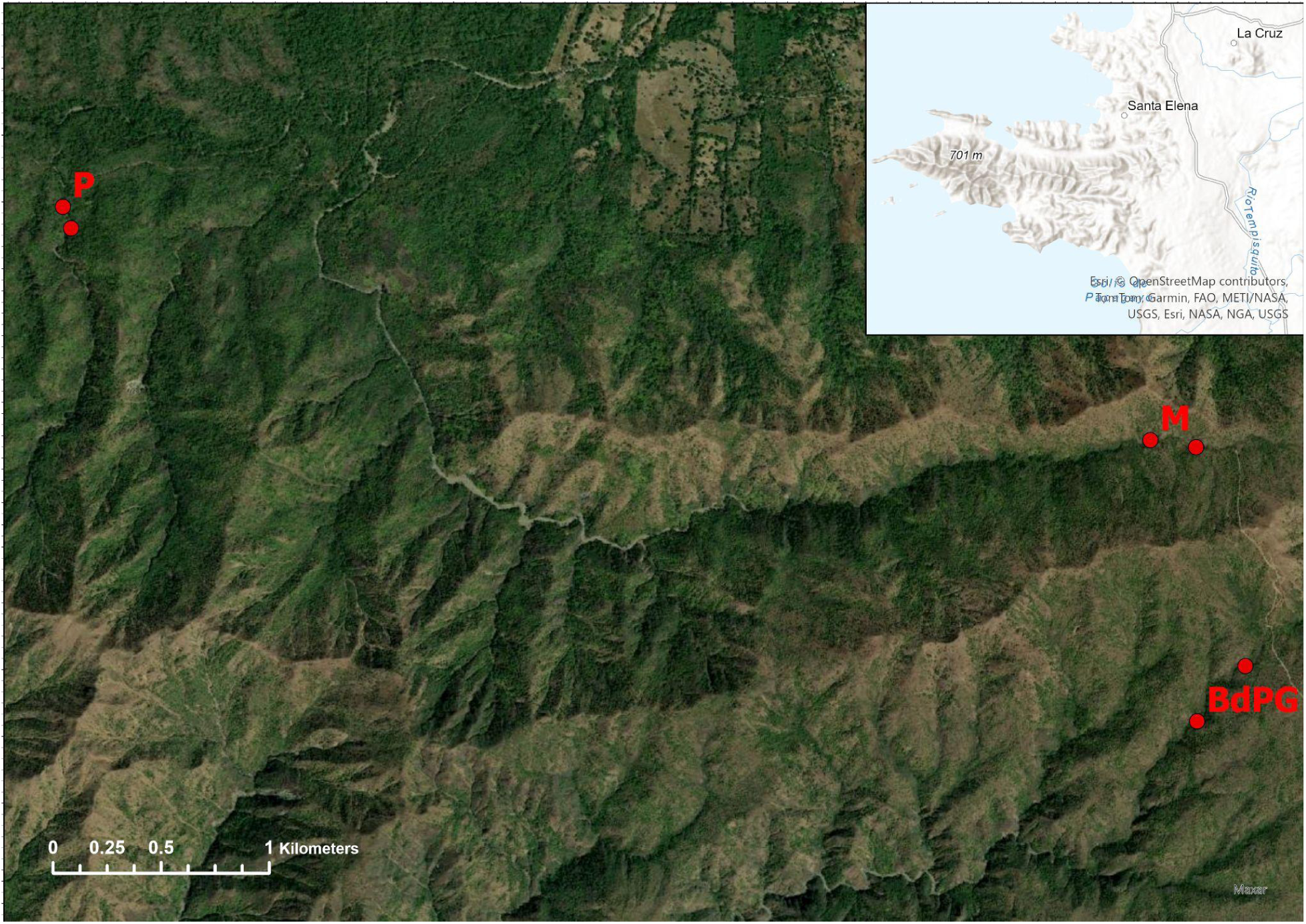
Sampling locations in northwestern Costa Rica. The main panel shows high-resolution satellite imagery of the study area, with sites labeled as M (Murciélago), P (Pedregal), and BdPG (Brazo del Río Potrero Grande). Red circles indicate start and finish points of transects where amphibian skin swabs were collected for microbial community characterization in Santa Elena Peninsula. Base imagery: Esri World Imagery basemap (© Esri, Maxar, Earthstar Geographics, USDA, USGS, AeroGRID, IGN, and the GIS User Community).

### Translocation and Radio Tracking

On March 10, 2023, selected individuals of *Craugastor ranoides* (>5g mass) were translocated to Maritza Biological Station, situated at the foothills of Volcán Orosí, an area where the species historically occurred. Records from nearby Volcán Cacao document its presence before the mass declines of the late 1980s, likely caused by chytridiomycosis outbreaks (Puschendorf et al., 2019). This site was selected because it lies just beyond the climatic envelope from which amphibian populations largely disappeared during the epidemic (Figure 2). To monitor short-term adaptation and habitat use following translocation, we employed radio tracking—the only feasible method for assessing individual movement and persistence at fine scales in this context. Prior to translocation, we conducted a pilot study in the dry forest to validate radio tracking techniques and assess frog behavior in their original habitat (Wakeham et al., unpublished), ensuring methods were appropriate and functional under local conditions. Only frogs weighing more than 5 grams were considered suitable for radio tracking, ensuring the transmitter weight did not exceed 10% of the individual’s body mass, in line with ethical guidelines for amphibian telemetry

**Figure 2.**
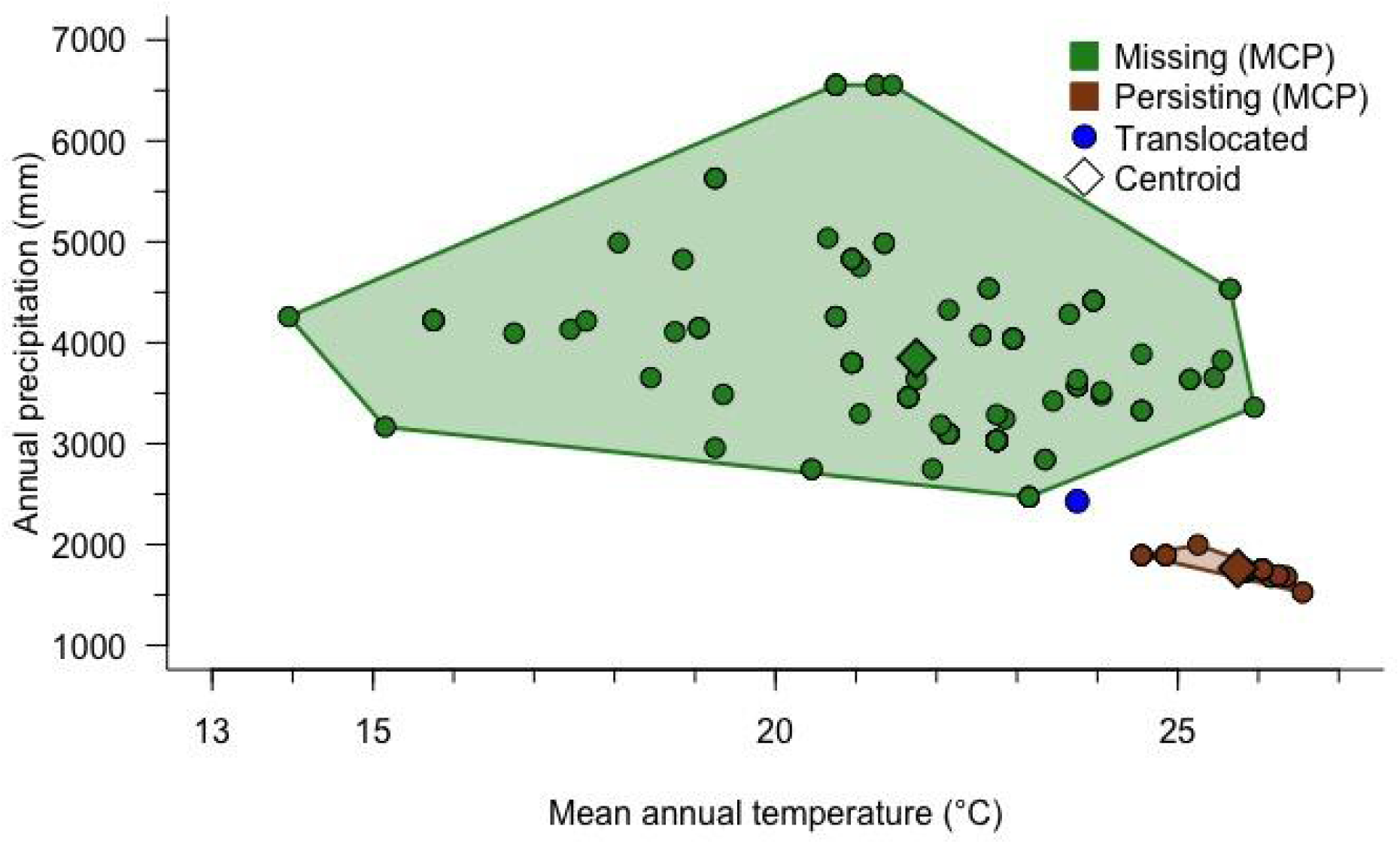
Climate space occupied by *Craugastor ranoides* presence sites across Costa Rica. Points represent sampling localities classified as persisting (current; brown circles), missing (historic; green circles), and translocated (blue circle). Semi-transparent polygons depict minimum convex polygons (MCPs) encompassing the persisting and missing sites, with solid rhomboids indicating group centroids. Climate data were extracted from CHELSA v1 (Karger et al., 2017) and compiled from historic and current records from the Museum of Zoology, University of Costa Rica and RP’s unpublished data.

(Richards et al., 1994; Muths, 2003). Frogs were weighed using Pesola® Light-Line Precision Spring Scales (Pesola AG, Baar, Switzerland) and measured for snout-vent length (SVL) using Proster® digital calipers (Model VC4102, Shenzhen, China) both prior to and following transport.

After sampling, each frog was placed in an individual sterile polyethylene bag (Whirl-Pak®, Nasco, USA) and transported in a Coleman® 30-Quart Performance Cooler (Newell Brands, USA) directly to Maritza Biological Station—approximately a one-hour drive from the capture location.

Upon arrival, frogs were reweighed and re-measured, and fitted with LB-2X radio transmitters (0.27 g, Holohil Systems Ltd., Ontario, Canada) using waist-belt harnesses constructed from silicone tubing, designed to minimize restriction of movement.

Radio tracking was conducted daily using a Lotek Biotracker VHF receiver (138–174 MHz) paired with a Lotek LiteFlex 3-element Yagi directional antenna, both purchased from Lotek Wireless Inc. (Newmarket, Ontario, Canada). This allowed for consistent post-release monitoring of frog movements, habitat use, and survival to assess adaptation to the new environment and evaluate the effectiveness of the translocation strategy.

### Sequencing and Bioinformatics

We extracted DNA from swabs using Zymbiomics Miniprep kits (Zymo, USA) using the standard kit protocol after removing ethanol from samples using a SpeedVac. We quantified DNA concentration using a Qubit 4 Fluorometer and sent normalised DNA extracts to the Exeter sequencing service where they prepared sequencing libraries by amplifying the V4 region of the 16S rRNA gene using primers 515F and 806R. We sequenced all libraries on an Illumina NovaSeq using 250bp PE reads.

We cleaned and quality-controlled raw sequences using *dada2* (Callahan et al., 2016) to identify Amplicon Sequence Variants (ASVs) and assigned taxonomy to ASVs using the SILVA reference database (v138.1;(Callahan, 2024; Quast et al., 2012). We removed potential contaminant sequences using *decontam* (Davis et al., 2018) prior to downstream analysis.

### Statistical Analysis

*We* used *phyloseq* (McMurdie and Holmes, 2013) and *vegan* (Oksanen et al., 2018) for downstream processing of cleaned microbiome data, alongside *ggplot2* (Wickham, 2016), *ggordiplots* (Quensen et al., 2024) and *cowplot* (Wilke, 2024) for visualisation of microbiome data. We used *vegan* for PERMANOVA and ordinations, *brms* (Bürkner, 2021, 2018) to fit Bayesian generalized linear models, and *gllvm* (Niku et al., 2025, 2019) to fit General Linear Latent Variable models (GLLVMs). We used *ComplexUpset* (Krassowski et al., 2022; Lex et al., 2014) to visualise shared and unique ASVs by site ID, and *NetCoMi* (Peschel, 2022) and *SpiecEasi* (Kurtz et al., 2024) to perform network analyses by quantifying (dis)similarities among both samples and ASV abundance data.

Final post-QC library sizes ranged from 15,189 - 176,202 reads per sample across both the Spatial and Translocation samples (mean 102,663). We used species rarefaction curves fitted in *vegan* to assess coverage of community composition, where all curves plateaued at <15,000 reads suggesting we had adequately sampled microbiome communities.

#### Spatial Dataset

To examine differences in alpha diversity (richness), we fitted a Negative Binomial GLM in *brms* with number of ASVs as the response, and SVL (mm) and site ID as predictors, where site is a 3-level factor describing key sampling sites: Brazo del Poterero Grande (BdPG), Murcielago (M), & Pedregal (P); Fig. 2). To examine differences in beta diversity, we performed PCA ordination on CLR-transformed relative abundances to account for the compositionality of sequencing data (Gloor et al., 2017). To model drivers of beta diversity, we fitted a PERMANOVA on CLR-transformed abundances with site (3 level factor) and SVL as predictors, specifying 999 permutations and a Euclidean distance metric.

To examine the effect of site and SVL on the relative abundance of individual microbial genera, we fitted a GLLVM specifying k-2 latent variables. We used only the top 50 most abundant bacterial genera for this analysis. We built a sample similarity network based on Aitchison distance using *SpiecEasi* and *NetComi* using CLR as a normalisation method, K-Nearest-Neighbour to account for sparsification, followed by greedy clustering.

#### Translocation Dataset

We visualised changes in beta diversity using a PCA ordination on CLR-transformed abundances, and a heatmap of the top 50 most abundant ASVs using *pheatmap* (Kolde, 2019). We fitted a PERMANOVA with timepoint (Pre- and Post-Translocation and ID as fixed effects, specifying a constrained permutation structure by ID to control for repeated measures. We built a sample similarity network from the raw ASV abundances using the same specifications as the spatial dataset. To investigate differences in individual ASVs associated with translocation, we fitted a GLLVM with time point as a fixed effect and individual ID as a random effect, with 2 latent variables.

## RESULTS

### Spatial Patterns in Skin Bacterial Community Structure

We generated microbiome community data from a total of 48 *C. ranoides* individuals sampled across three discrete populations. We found limited evidence for variation among populations in average bacterial richness (alpha diversity), instead finding substantial within-population variation (Fig. 3A). A model containing fixed effects of population ID and SVL was an inferior fit to an intercept only model (null model LOO-IC = 560.2, full model LOO-IC = 561.4). There was also no evidence for an effect of pH on patterns of bacterial richness (cor = −0.15, p=0.3, Fig. S1).

**Figure 3.**
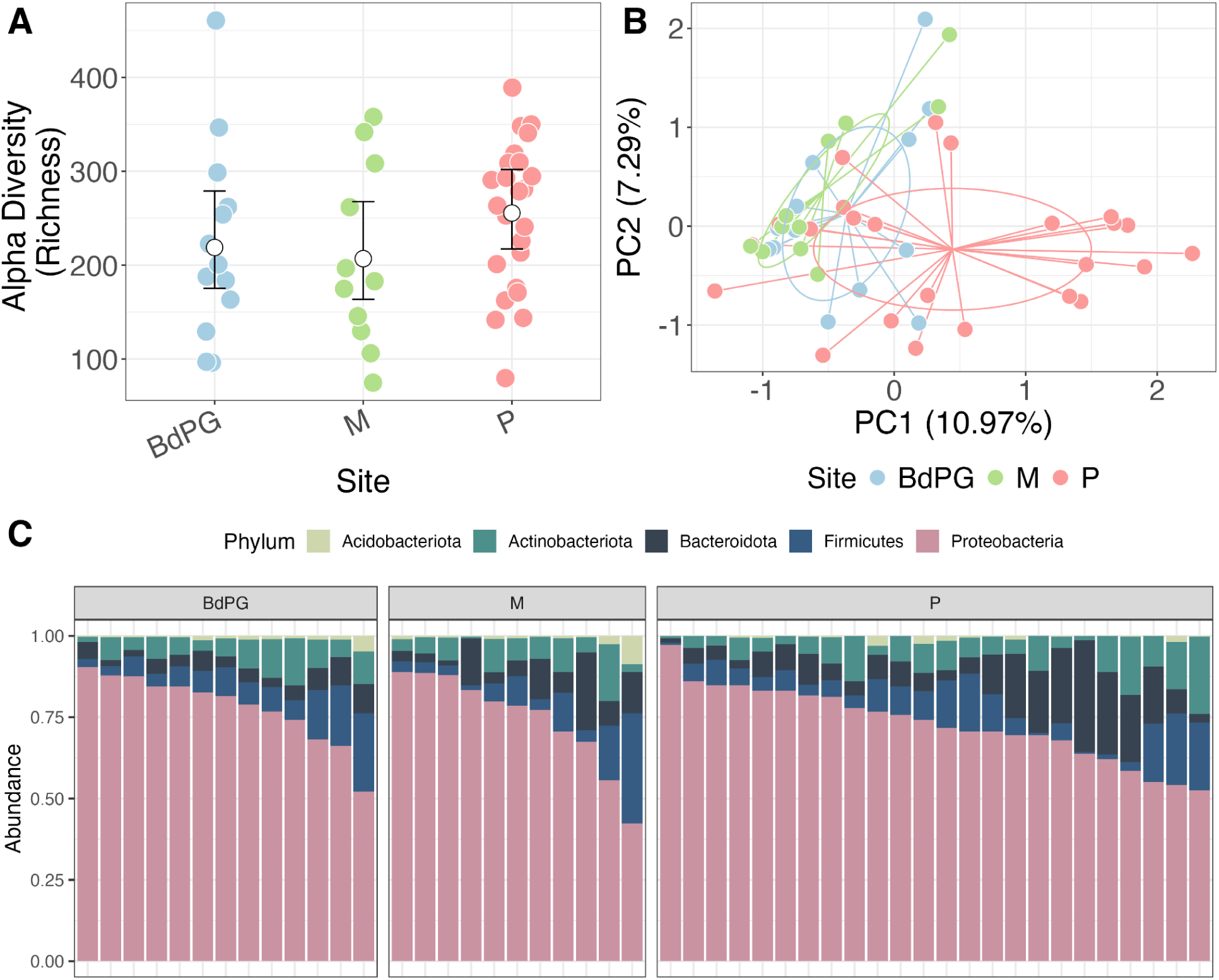
Bacterial community composition variation in space. A) No evidence of variation in mean bacterial richness among sites. Coloured points are raw data means. White points are posterior model means from a Bayesian GLM. Bars represent 95% credible intervals. B) PCA of Centred log-ratio bacterial community composition, coloured by site. Ellipses represent 95% confidence intervals of the mean and lines connect raw data (points) to site-specific centroids (weighted averages). C) Stacked bar plots of the relative abundance of the top 5 bacterial Phyla, ordered by proportion of Proteobacteria and site ID (facets).

We did detect significant variation among populations in bacterial community composition (beta diversity; PERMANOVA effect of site (F_2,40_ = 1.68, p=0.001, r^2^=7%; (Fig. 3B,C). This effect seemed to be largely driven by divergence of several Pedregal individuals from the other sampling locations (Fig 3B). Though all sites shared a large number of ASVs (Fig. 4A), Pedregal has the highest number of unique ASVs that explain in part its dissimilarity to samples from other sites (Figs. 3B; 4A). There was also substantial variation in community composition among individuals *within* sites; at the Phylum level some individuals were dominated by Proteobacteria whilst others had comparably more members from Firmicutes and Acidobacteria. We found no effect of SVL on beta diversity (PERMANOVA F_1,43_ = 1.19, p=0.13, r^2^=2.5%).

**Figure 4.**
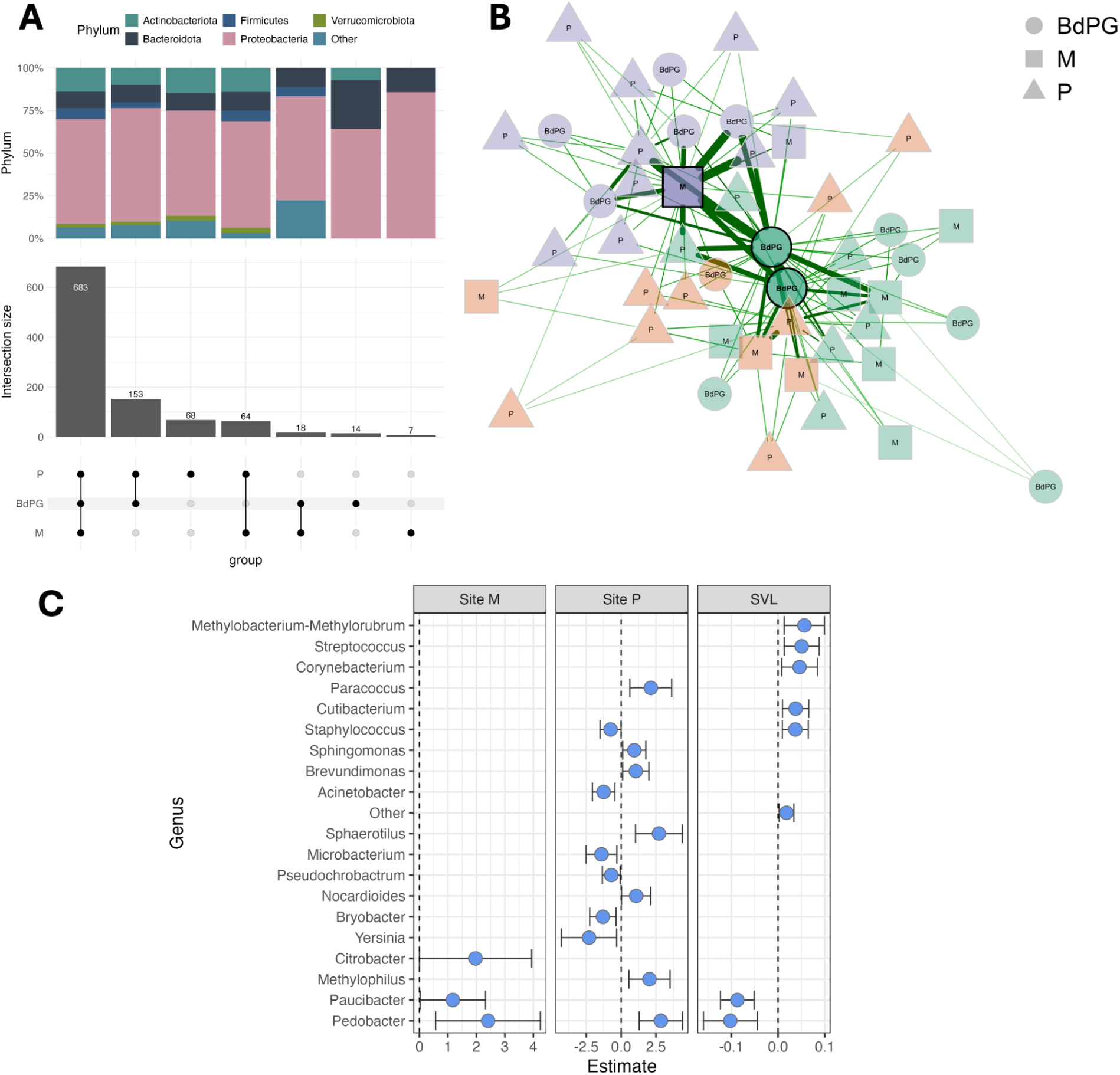
Difference in skin bacterial composition by Site and SVL. (A) Upset plot showing overlapping and unique ASVs per site. Upper stacked barplot shows taxonomic composition at the Phylum level of ASVs for each group. (B) Sample similarity network for spatial study. Each sample is a node; connections between nodes indicate stronger covariance in ASV abundances between that pair of samples, with thicker edges showing higher relative similarity. Highlighted nodes are ‘hub’ nodes with high eigenvector centrality - connected to nodes that themselves are well connected. (C) Output of a GLLVM showing significantly different ASVs in sites P and M (compared to BdPG as the intercept), and SVL.

Building a sample similarity network did not reveal any systematic clustering by population (Fig. 4B). Individuals from BdPG and Murcielago were identified as ‘hub’ nodes with high centrality in the network, and strong similarity to other samples. Though the network analysis identified 3 distinct statistical clusters, samples from all 3 main populations were divided among them. However, Pedregal samples were mostly found on the periphery of the network, with lower similarities to fewer samples, on average (Fig. 4B). Collectively these data support the analyses of beta diversity showing marked within-population heterogeneity in composition alongside a large core microbiome of shared ASVs (Fig. 4A) -both of which are expected to disrupt the emergence of strong separation of samples by population of origin.

Using a GLLVM we found several changes at the genus level linked to site ID (Fig. 4C). Using site BdPG as the intercept, frogs from Pedregal showed increases in the relative abundance of the genera *Sphingomonas, Pedobacter, Paracoccus,* and *Brevundimonas*, and decreases in multiple genera including *Yersinia* and *Microbacterium* (Fig 4C). Site M had comparably fewer differentially abundant ASVs, supporting the patterns seen in whole-community ordination (Fig. 3B). Larger frogs (greater SVL) had higher relative abundances of *Streptococcus, Staphylococcus* and *Corynebacterium*, and lower abundances of *Pedobacter* and *Paucibacter* (Fig. 4C).

### Temporal Dynamics Linked to Translocation

Of the 14 *Craugastor ranoides* individuals translocated, all contributed microbiome samples at the pre-translocation timepoint, but only nine were recaptured and sampled again post-translocation, as microbiome swabbing was timed to coincide with transmitter removal, and some transmitters were not reachable at this point. Of the five frogs not recaptured, one individual’s transmitter was detected under a large boulder approximately five meters from the stream, suggesting predation, possibly by a snake. The remaining four were intermittently detected earlier in the study but could not be located at the final time point, because they could not be accessed (frog hiding and inaccessible), or transmitter failure during the course of the radio tracking period. Among these frogs sampled before and after transmitter removal, body weight changes were variable: five frogs gained weight, three lost weight, and one remained unchanged, with a mean weight change of +0.39 g (Fig. 5A).

**Figure 5.**
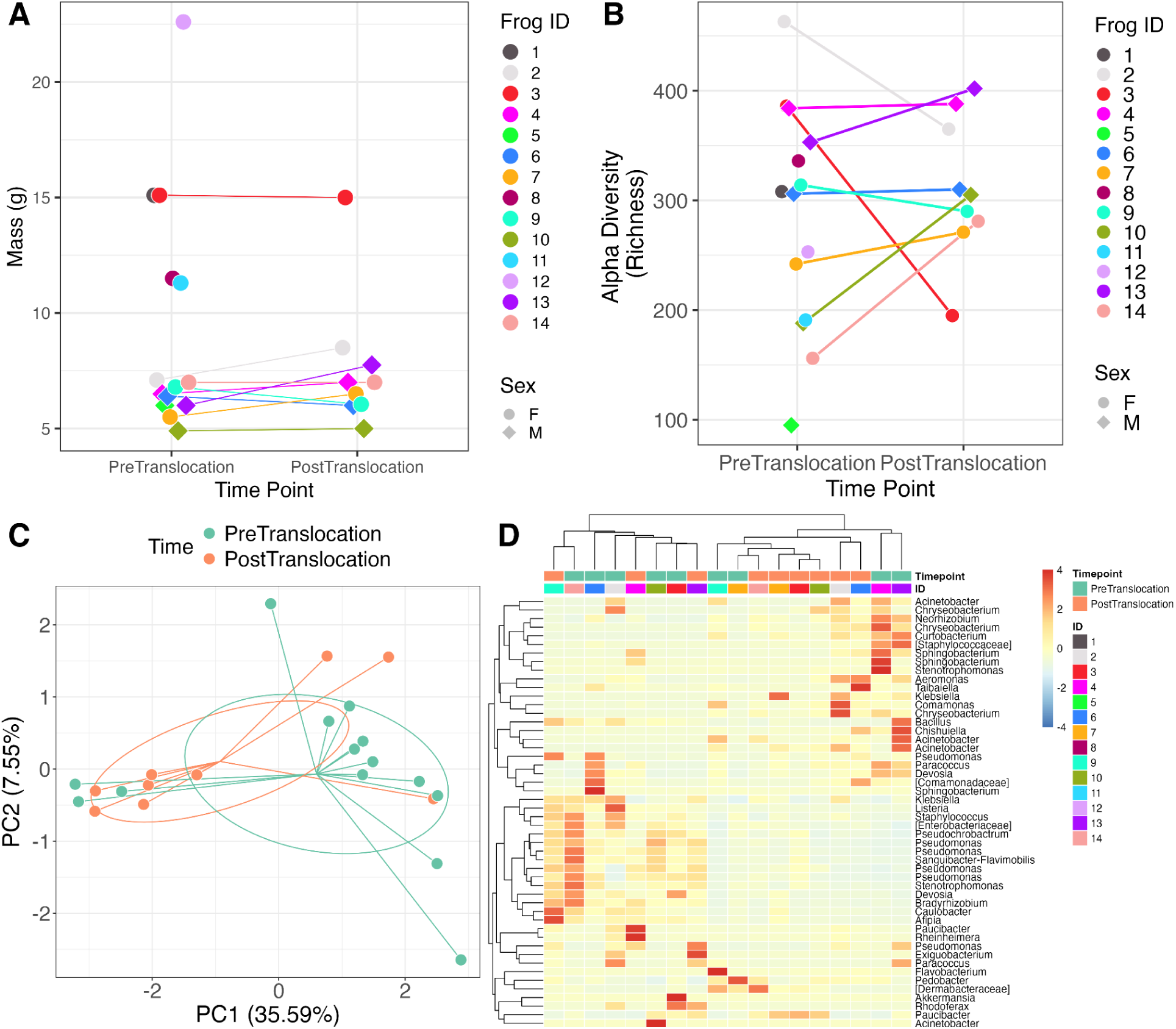
Temporal microbiome dynamics following translocation. A) Changes in body mass at the individual level. Lines link repeat sampled frogs caught 3-4 days after translocation. Some individuals were not recaptured and so are not connected to other points. B) Changes in bacterial richness (alpha diversity). Lines link repeat sampled frogs caught 3-4 days after translocation. Some individuals were not recaptured and so are not connected to other points. B) Ordination of CLR-transformed bacterial communities by translocation. Lines join points to group centroids and ellipses are 95% confidence intervals of the centroid location. D) Heatmap of relative abundances of the top 50 most abundant ASVs, clustered by similarity.

We found that no consistent change in alpha diversity linked to translocation; individual ‘reaction norms’ pre- and post-translocation exhibited increases, decreases, and no change in richness (Fig. 5B). These patterns were supported by ordinations of beta diversity that showed no consistent shift in location following translocation composition (Fig. 5C; PERMANOVA effect of time (translocation) F_1,8_=0.67, p=0.87; Procrustes rotation, cor=0.49, p = 0.27). There was a strong signal of individual ID in bacterial community composition (r^2^=0.46), but this effect was not significant (F_13,8_ = 0.61; p=0.99).

We found no clear clustering in a heatmap of samples by timepoint (pre- vs post translocation) or individual ID (Fig. 5D). However, some sets of samples did show similar compositional profiles and clustered together, including samples from different individuals from different timepoints (Fig 5D). In the sample similarity network, most of the pre-translocation samples belonged to the statistical cluster (Fig. 6A), but strong connections between pairs of samples from the same individual were rare, suggesting strong within-individual divergence post-translocation. That some individuals maintained similar levels of richness either side of translocation (Fig. 5B) but remain poorly connected in the network (Fig. 6A) is indicative of significant turnover in membership of their bacterial microbiota.

**Figure 6.**
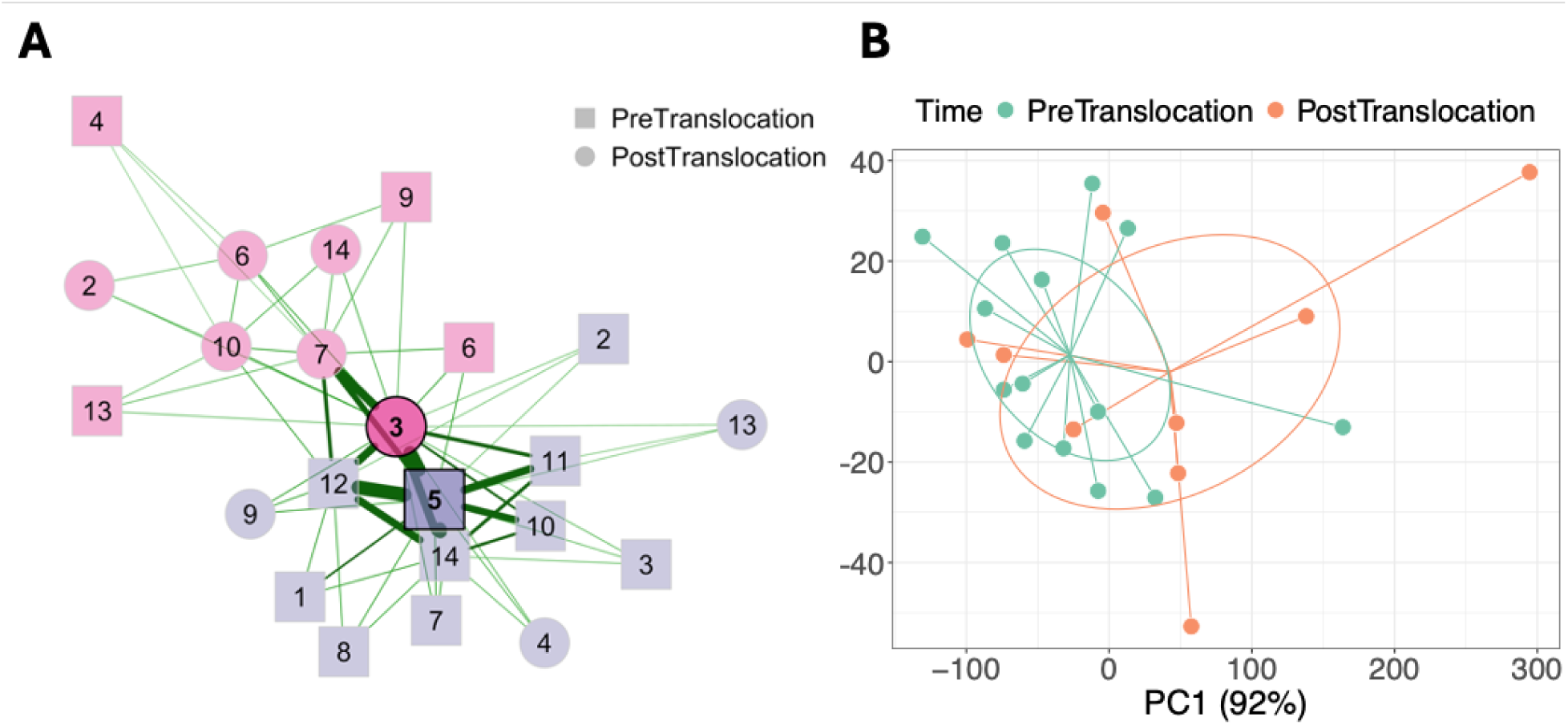
Changes in microbiome composition and predicted function associated with translocation. (A) Sample similarity network for translocated individuals.Shape indicates time point (pre- vs post-translocation. Colour denotes statistical cluster, and edge weight indicates sample similarity. The two highlighted ‘hub’ nodes have the highest eigenvector centrality. Some repeat-sampled individuals had no network connections (no edges with other samples) and so were removed from this network. (B) NMDS ordination of changes in predicted microbiome function associated with translocation.

Using a GLLVM we investigated changes in the relative abundance of specific ASVs linked to translocation whilst controlling for repeated measures on individuals. We found that after translocation, there were increases in ASVs from the genus *Curvibacter,* and decreases in *Devosia*. Using the 16S sequences to predict the functional profiles of skin-associated bacterial communities, we found no consistent shifts in functional profile associated with translocation (PERMANOVA; F_1,8_ = 0.79, p = 0.78, r^2^ = 3.8%; Fig 6B). There was a strong signal of individual ID in predicted functional profile (r^2^ = 57.2%), though the effect of ID was not significant (F_13,8_ = 0.9, p=0.24).

## DISCUSSION

Translocations are an important tool for species management and conservation, especially for amphibians in the context of mitigating disease-driven biodiversity declines (Scheele et al., 2021). Performing translocations is logistically challenging, and translocation success is dependent on a broad range of biotic and abiotic factors, including host behaviour and quality of the release habitat (Berger-Tal et al., 2020; Hossack et al., 2022a) or persistence of lethal pathogens (Scheele et al., 2021). However, only recently has the potential importance of host-associated microbial communities as critical agents of translocation success been recognised (e.g. Chong et al., 2019; Dallas and Warne, 2023b; West et al., 2019b; Worsley et al., 2024). As a result, few studies have integrated metrics of microbiome composition into decisions of which populations or individuals to translocate, nor the impact of translocation on subsequent microbiome dynamics (but see Worsley et al., 2024a). Here we have assessed the extent of among-population variation in skin bacterial communities of the critically endangered frog *Craugastor ranoides*, and assessed impact of translocation on microbiome dynamics at the individual level. We found substantial within-population variation in the skin microbiota, and no evidence of a consistent directional shift in bacterial community composition associated with translocation. Instead we detected strong ‘individuality’ of microbial dynamics; some individuals increased the richness of their microbiomes, whilst others exhibited a stable alpha diversity with marked turnover in the identity of bacterial membership.

### Limited Among-Population Variation in Skin Bacterial Communities

We found limited among-population variation in bacterial community richness, and only weak differentiation among populations in beta diversity. Instead, all three populations demonstrated equal or greater within-rather than among-population variation in traits of bacterial community composition. Limited among-population divergence in alpha diversity have been observed in the gut microbiomes of passerine birds (Liukkonen et al., 2024), but for many species the general pattern is that spatial structure is associated with among-population divergence in microbiota similarity (reviewed in Couch and Epps, 2022), especially for amphibian skin microbiomes (e.g. Griffiths et al., 2018; Kueneman et al., 2014) that are more environmentally labile (Harrison et al., 2019). In our study, the water courses at all three sampling sites had similar pH values in the range of 7.8-8.3, which may constrain variation in skin bacterial communities (Muletz Wolz et al., 2018). Despite this, Pedregal showed the largest variability in skin microbiota composition, with the highest number of unique bacterial ASVs and greatest dispersion in microbial community composition in ordination space.This elevated variability may reflect unique environmental conditions at Pedregal. Unlike the other two sites, Pedregal contains active serpentinizing springs, features where ultramafic rocks react with water, producing high-pH fluids (up to ∼11), elevated calcium concentrations, and methane-rich emissions (Sánchez-Murillo et al., 2014). These geochemical gradients are known to support dense microbial biofilms and may influence the availability and composition of environmental bacteria available for skin colonization. The alkaline, chemically distinct conditions created by serpentinization may thus contribute to the increased dispersion and uniqueness of skin bacterial communities observed at this site.

A key unanswered question is what drives the observed *within*-population variation in skin microbiome. We found no clear relationship between SVL and *average* community composition, suggesting no consistent wholesale change in microbiome structure as hosts grow and age (Campbell et al., 2019). Using GLLVMs, we did detect subtle shifts in the relative abundance of particular bacterial genera when controlling for differences among sites. These included increases in the relative abundance of *Streptococcus* and *Staphylococcus* and *Corynebacterium*, all of which are common residents of the amphibian (Niederle et al., 2019) and wider vertebrate skin microbiome (Ross et al., 2019). We also detected decreases in the relative abundance of *Paucibacter* and *Pedobacter* with increasing SVL. *Paucibacter* is common in freshwater environments in the tropics, and has been found in both the nest microbiota of foam-nesting frogs (Monteiro et al., 2023) and skin bacteriome of cascade frogs (Roth et al., 2013). Changes in the relative abundance of these ASVs in larger frogs are likely to reflect differences in the manner in which frogs of different body size interact with the environment (Kelleher et al., 2017), a key determinant of the amphibian skin microbiome (Harrison et al., 2019). For example, larger frogs may reduce time spent within or interacting directly with the water course, and so undergo a corresponding decrease in freshwater-associated bacteria. Observations during radio-tracking suggested that females were often more likely to leave the stream and move into adjacent terrestrial habitats, whereas males appeared more territorial and tended to remain closer to the water course. This pattern is consistent with field studies of other stream-breeding Craugastor species, where females were more difficult to capture and exhibited more extensive movement across stream habitats (Zumbado-Ulate et al., 2011). The functional consequences of these changes remain unknown; genera such as *Paucibacter* can hydrolyze amino acids and so provide a nutritive substrate for other community members (Rapala et al., 2005). Decreases in the prevalence of microbes performing these roles may be expected to change the fundamental dynamics of interactions among members of the microbiome.

### Temporal Dynamics of The Microbiome Linked to Translocation

We predicted that translocation would be associated with a consistent turnover in microbiome composition, and expected that individuals post-translocation would have more similar microbiomes to one another than to their pre-translocation counterparts. Our data did not support this prediction; for some individuals we observed both gain and loss of bacterial community richness over time, whilst others showed stable patterns of richness but underwent significant changes in microbial community membership. There was also no evidence of strong individual stability of the microbiome, which would manifest as resistance to changes in community membership following translocation. We also found no clear signal of a shift in predicted functional repertoire of the microbiome either side of translocation. Microbiomes are highly dynamic (Marsh et al., 2024), and understanding the forces shaping that variability in microbiome structure and function remains a major research goal (Worsley et al., 2024b). Previous work on translocated Seychelles warblers found consistent reductions in alpha diversity associated with translocation (Worsley et al., 2024a), a pattern that contrasts with our data. Most prior research on the effect of translocation on the amphibian skin microbiome tends to focus on individuals that have first been captive-reared (Korpita et al., 2023; Kueneman et al., 2022), and so drawing comparisons to individuals moved from the wild is difficult given that captivity tends to shape microbiome in a unique manner (Kueneman et al., 2022). If anything these data serve to highlight just how little is known about the causes and consequences of host microbiome responses to translocation.

When controlling for individual ID, we found increases in the relative abundance of the genera *Curvibacter*, alongside decreases in *Devosia*. *Curvibacter* has been detected in the skin communities of several frog species (Roth et al., 2013). Shifts in relative abundance of these genera may simply reflect differential abundance in the environment of the translocation site, resulting in differential colonisation pressure on skin-associated communities (Harrison et al., 2019). Given how strong this environmental signal usually is for amphibian skin microbiotas, that we did not observe a convergence in bacterial community composition suggests other factors must be in effect. We suspect the most likely explanation for variation in the rate and extent of skin microbiota turnover we observed is individual variation in behaviour. Behaviour can shape the amphibian skin microbiome by changing how, when and for how long individual frogs interact with terrestrial and aquatic substrates (Xu et al., 2020). Though we monitored movement through nightly radio tracking in this study, our data are not at the resolution required for tracking fine-scale among-individual variation in behaviour that we can link to microbial profiles. Future work should combine such fine-scale data with contemporary environmental samples that permit quantification of the microbial communities capable of colonising the host in the novel translocation environment.

### Implications for *C. ranoides* and Wider Amphibian Conservation

Our data have several implications for our understanding of the importance of the microbiome in animal translocations. Recently these has been interest in the idea of optimizing the microbial communities of translocated hosts to match the destination environment (Seddon and Redford, 2025; Trevelline et al., 2019); this can be achieved either by ‘bioaugmentation’ where beneficial bacteria are installed into resident communities from *in vitro* cultures, or by choosing populations or individuals that are closely matched in microbial composition to the communities found at the destination. Bioaugmentation can be challenging because the benefit imparted by the novel microbe can change depending on context (e.g. pathogen genotype; Antwis and Harrison, 2018). For *C. ranoides*, all of our sampled populations had broadly similar bacterial profiles. This could be viewed as a benefit for translocation in that we are able to draw on multiple separate pools of individuals that *on average* will all possess similar ‘microbial toolkits’ when placed in the novel environment. These data underscore the importance of thorough surveys of wild donor populations prior to translocation (Seddon and Redford, 2025); depending on the system and nature of the destination, among-individual variability in the microbiome may be seen as either beneficial or suboptimal for the translocation program. The exact balance of these two depends on how plastic the host microbiome is following translocation; if microbiomes are relatively stable then one might value strong divergence in initial microbiome composition as a bet-hedging strategy against future, unknown challenges in the novel habitat, such as variable pathogen genotypes. Our data show that some changes in microbiota composition post-translocation might be beneficial for host health and /or resilience to challenges such as pathogenic infection. We detected increases in the abundance of bacterial genera known to possess *Bd*-inhibitory abilities, though explicit functional assays would be required to definitively determine how such bacteria may affect the outcome of host interactions with pathogens.

Beyond the microbiome, *C. ranoides* offers a compelling model for amphibian resilience. Previously, persistence of the species in the Santa Elena Peninsula, a hot, dry Pacific coastal region, was hypothesized to reflect a climatic refuge from Bd (Puschendorf et al., 2019, 2009, 2005; Zumbado-Ulate et al., 2014, 2014). However, our study sites within this area also intersect with zones of active serpentinization (Sánchez-Murillo et al., 2014), where ultramafic rock, water interactions generate alkaline waters and support distinct microbial films. Serpentinization alters water chemistry, often elevating pH above 9 and could directly or indirectly influence skin microbiota or the viability of the causal agent of chytridiomycosis, the fungus *Batrachochytrium dendrobatidis*.

These serpentinization-driven habitats may thus represent not only climatic but geochemical refugia for chytrid-sensitive species like *C. ranoides*. Understanding the interplay of local environmental chemistry and microbial dynamics is crucial for amphibian conservation strategies, particularly as climate change continues to shift the distribution of both hosts and pathogens.

Notably, 3.5 months after the experiment concluded, six *C. ranoides* individuals were found in the immediate vicinity where they had been tracked during the study, suggesting short-term survival and possibly site fidelity. While informal, this follow-up reinforces the possibility that translocated frogs can persist in suitable montane environments. Taken together, our findings highlight the value of incorporating microbiome dynamics and individual-level variation into amphibian translocation planning.

## Supporting information

Supplementary Plots

## DATA AVAILABILITY

Sequencing data for this project are stored in the Sequence Read Archive (SRA) under Project ID PRJNA1298236. Data and code to reproduce all analyses are available at https://github.com/xavharrison/RanoidesCostaRica24

## ETHICS AND PERMITS STATEMENT

This research was approved by the University of Costa Rica(UCR) and the Sistema de Areas de Conservacion de Costa Rica (SINAC). Permit resolutions CIBET-085-2022 and CBio-21-2022 were granted by UCR, and permits ACG-121-2021 and ACG-068-2022 were issued by. These permits comprehensively covered all aspects of the research, including ethical review, fieldwork, and molecular analyses, for the entire duration of the study.

## AUTHOR CONTRIBUTIONS

RP, XAH, KH, VW, AW, AN, HZ, MMC, JB, GA & RF designed and conducted the research, fieldwork and collected samples. KJM and XAH performed the laboratory work for DNA extraction. XAH performed bioinformatic & statistical analysis. XAH and RP wrote the manuscript, with comments from all authors.

