## Supplementary Plots for "Microbiome Dynamics During Translocation of the Critically-Endangered Frog *Craugastor ranoides*"

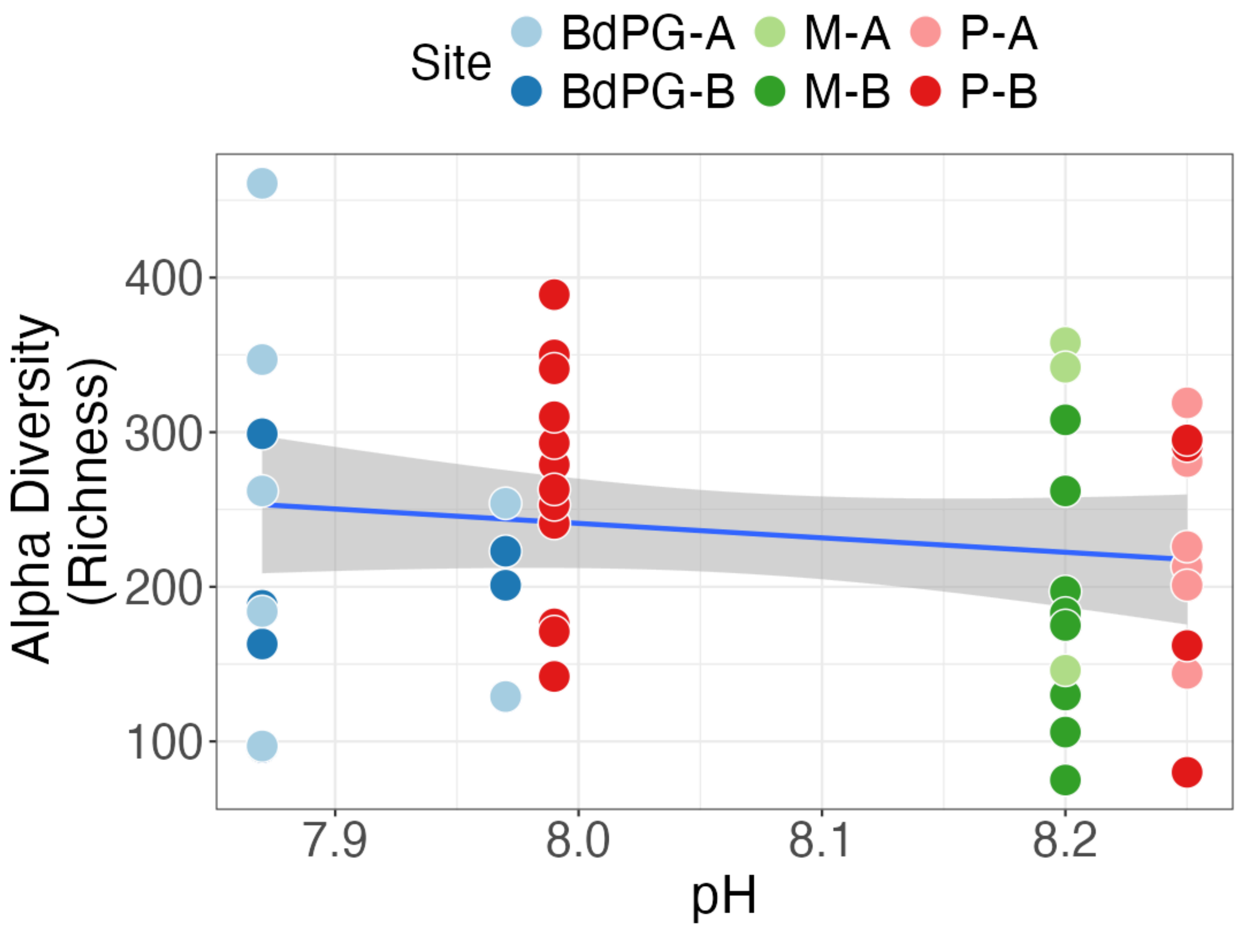


**FIgure S1.** Bacterial richness by pH of the sampling location

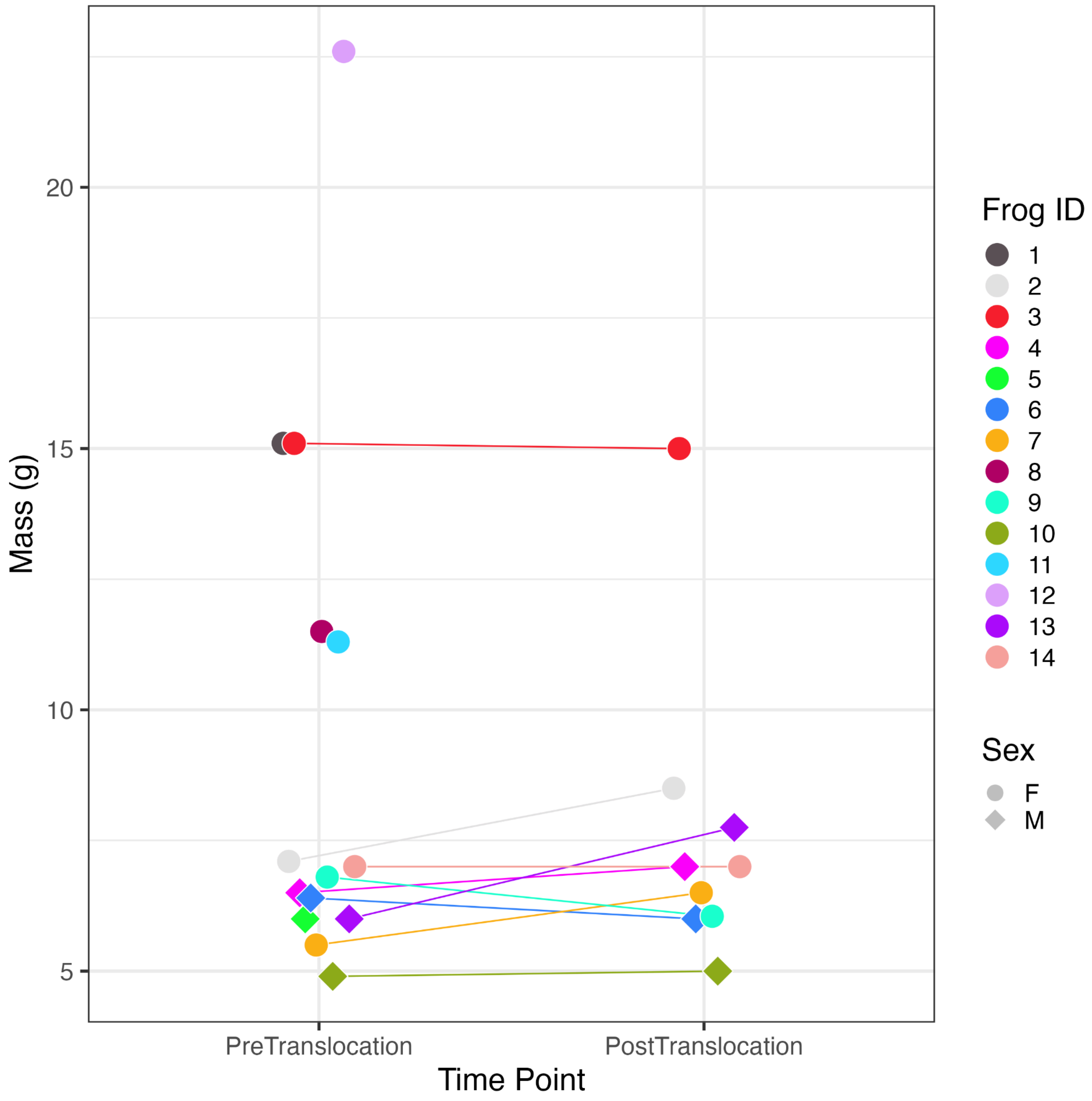


Figure S2. Body weight of Craugastor ranoides individuals before and after translocation Of 14 translocated frogs, 9 were recaptured. Each line connects the same frog pre- and post-translocation, shaped by sex (circles = Female, diamonds = Male).
